## Supplementary material for "Differential adhesion regulates neurite placement via a retrograde zippering mechanism": includes supplementary figures, legends, tables and Note

### Supplementary Fig. 1

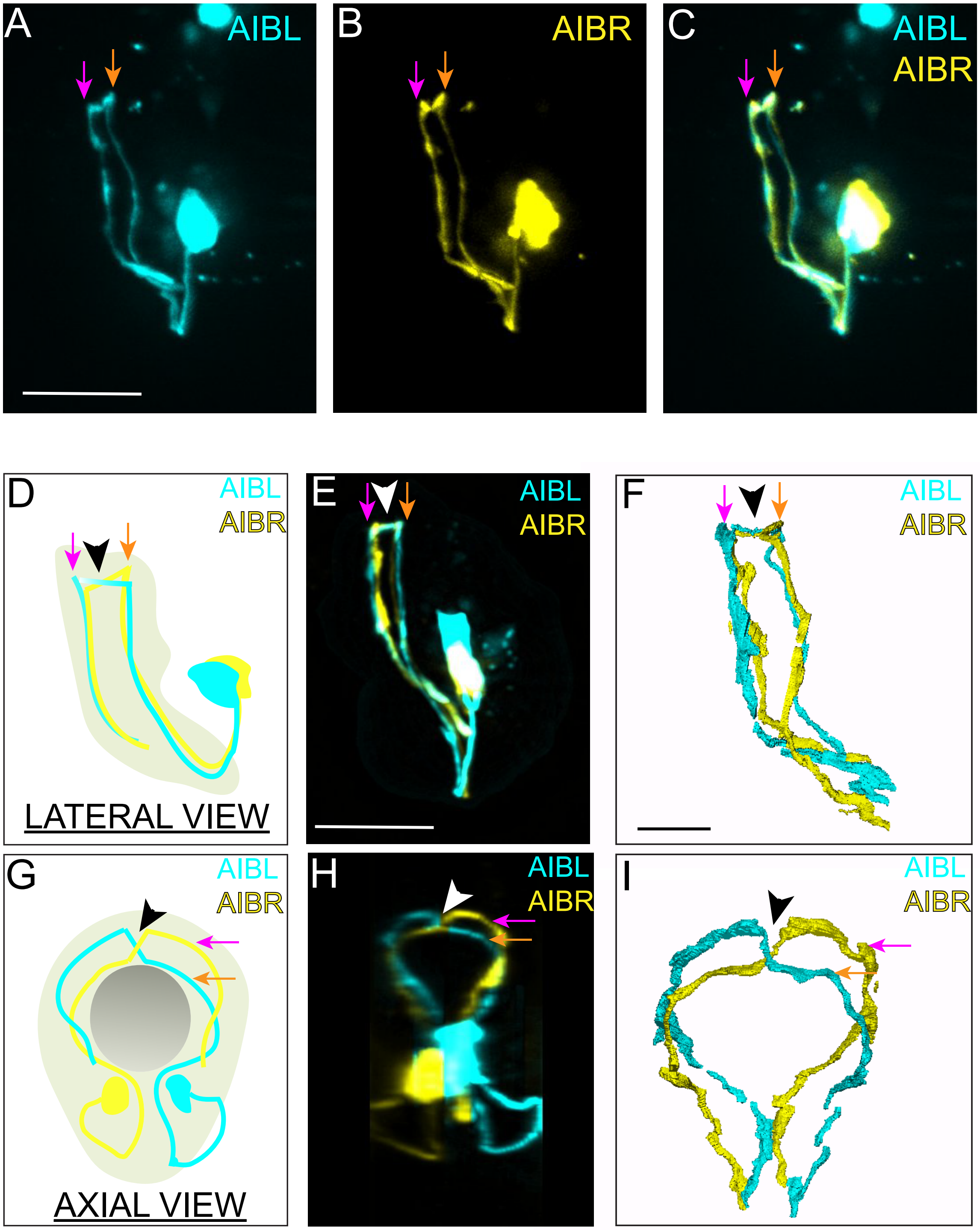

### Supplementary Fig. 2

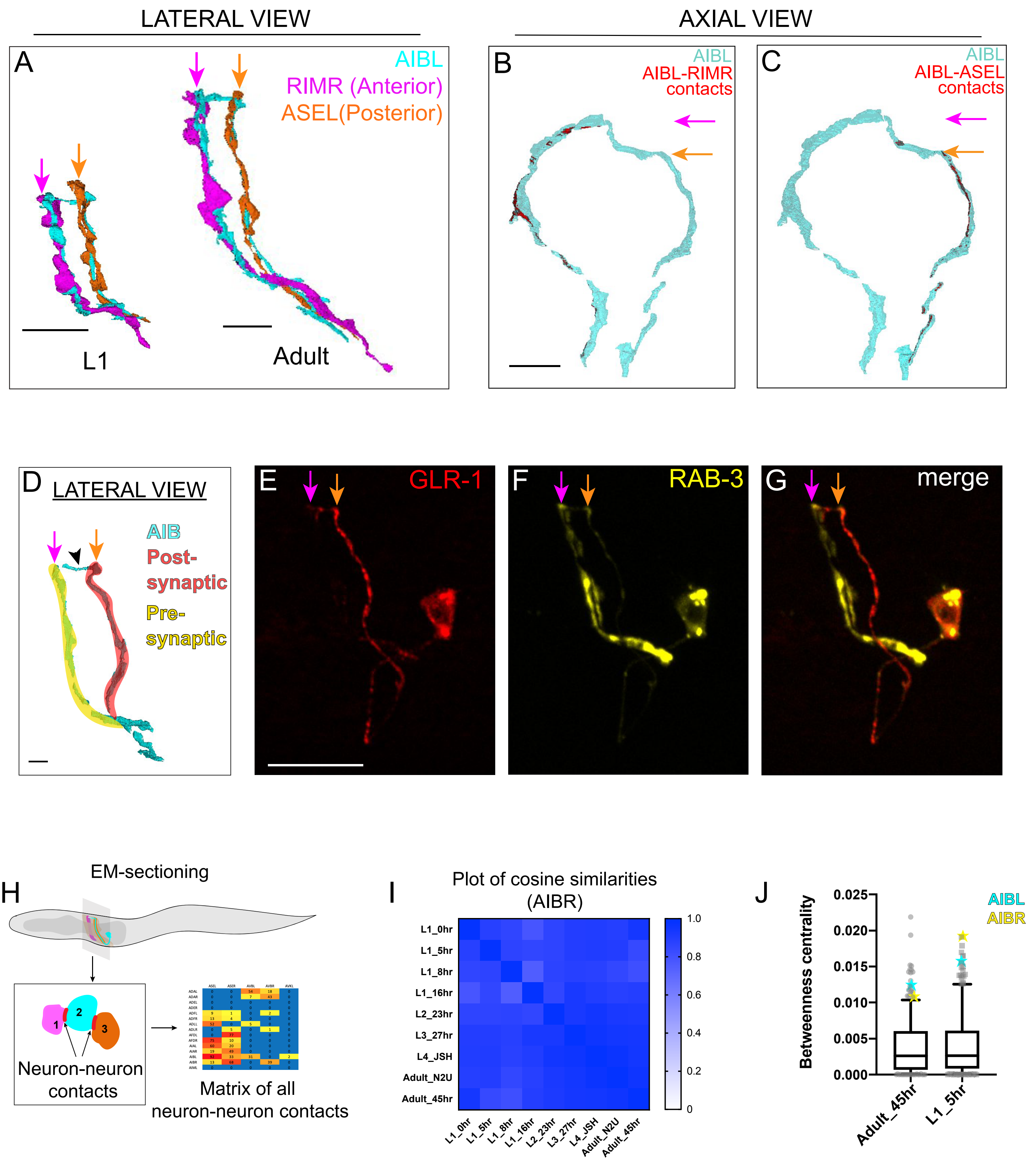

### Supplementary Fig. 3

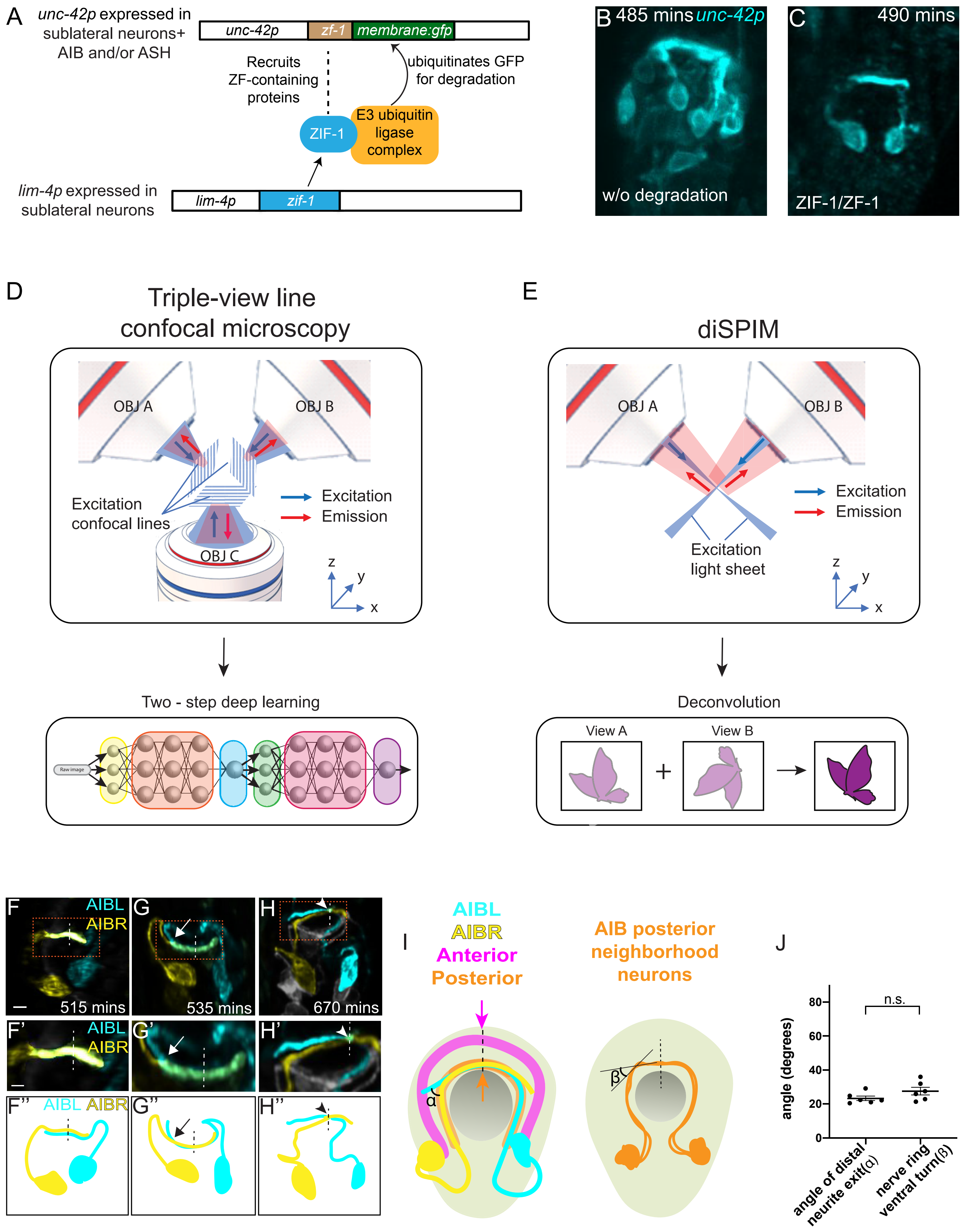

### Supplementary Fig. 4

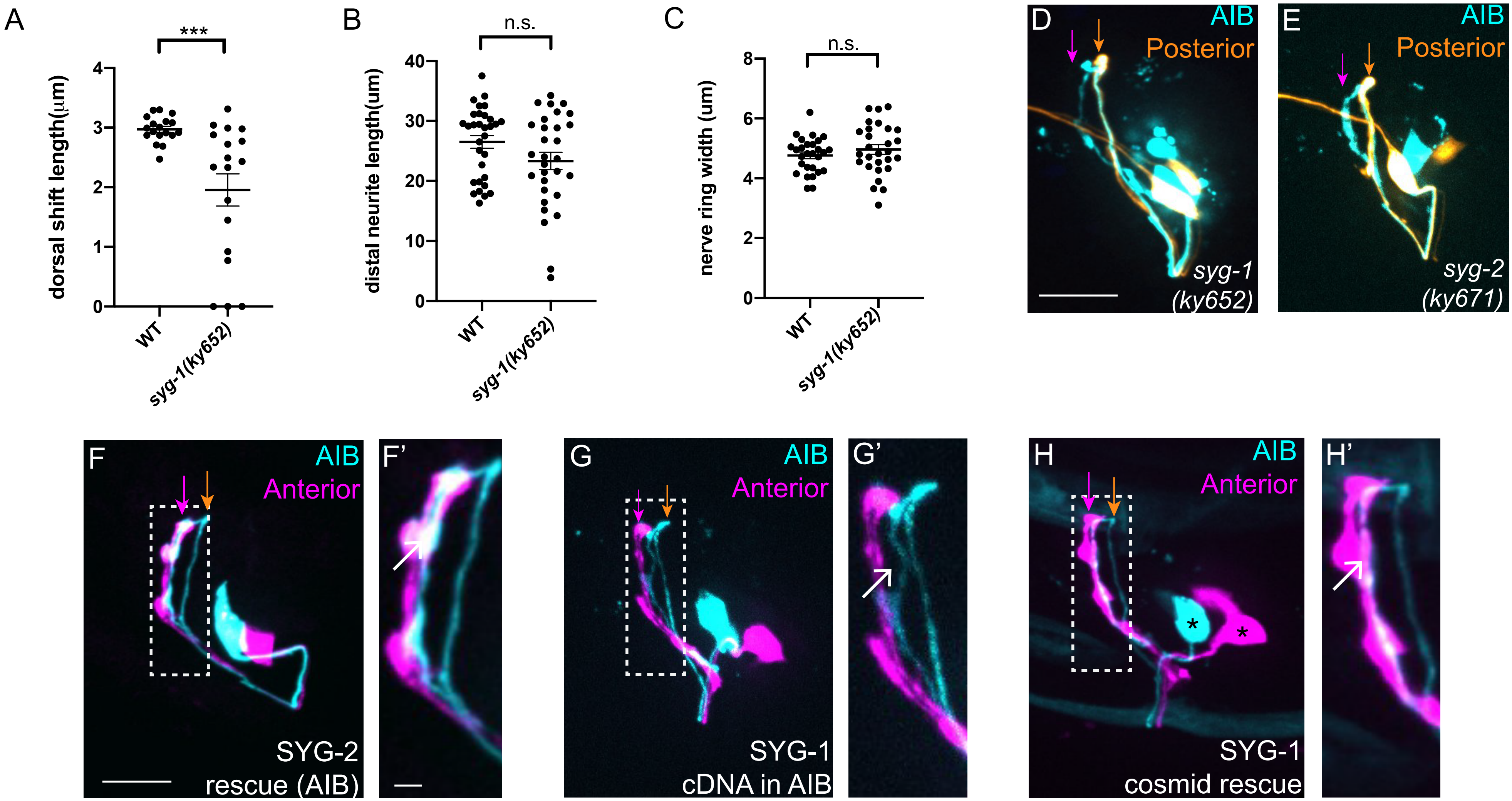

### Supplementary Fig. 5

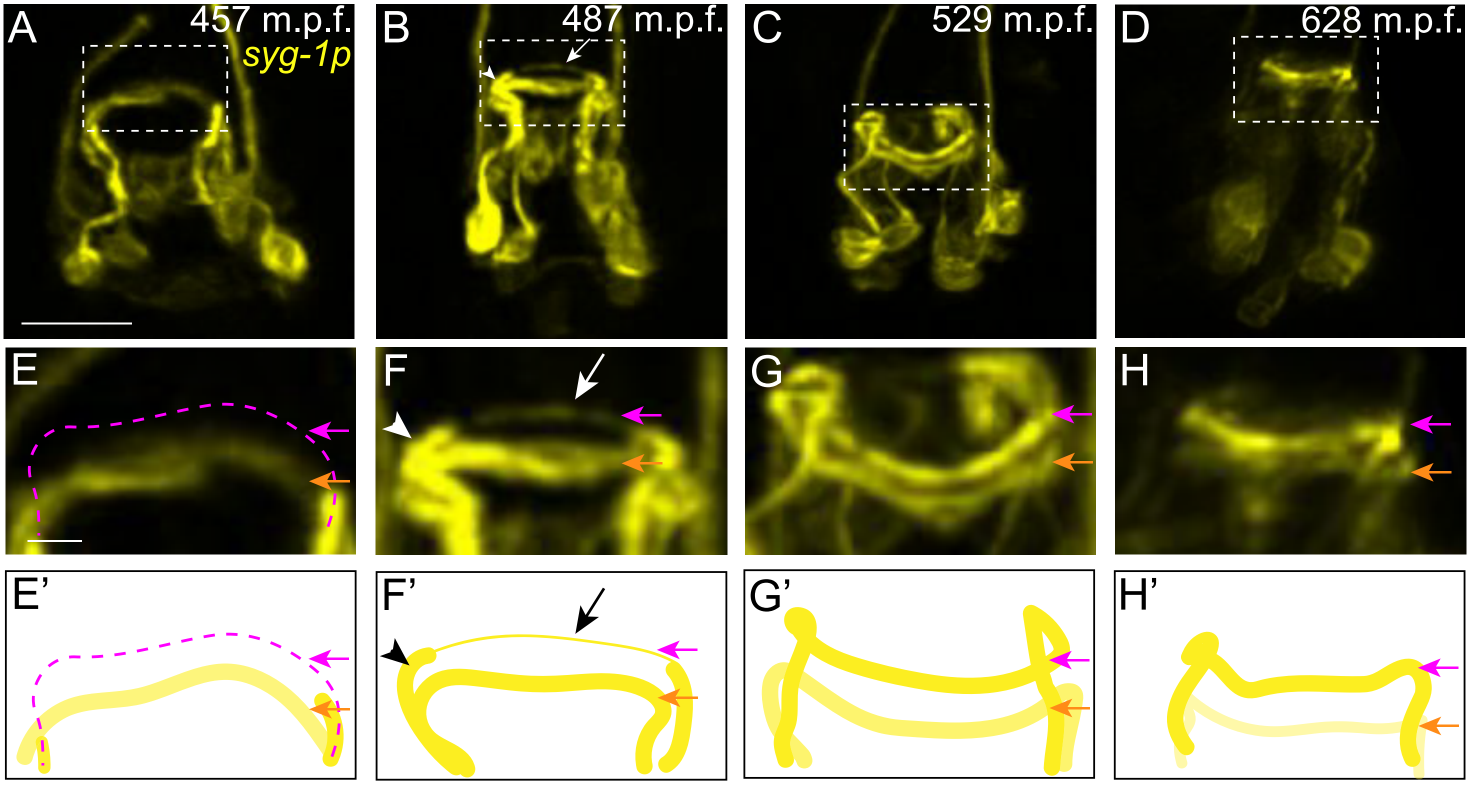

**Supplementary Fig. 6**

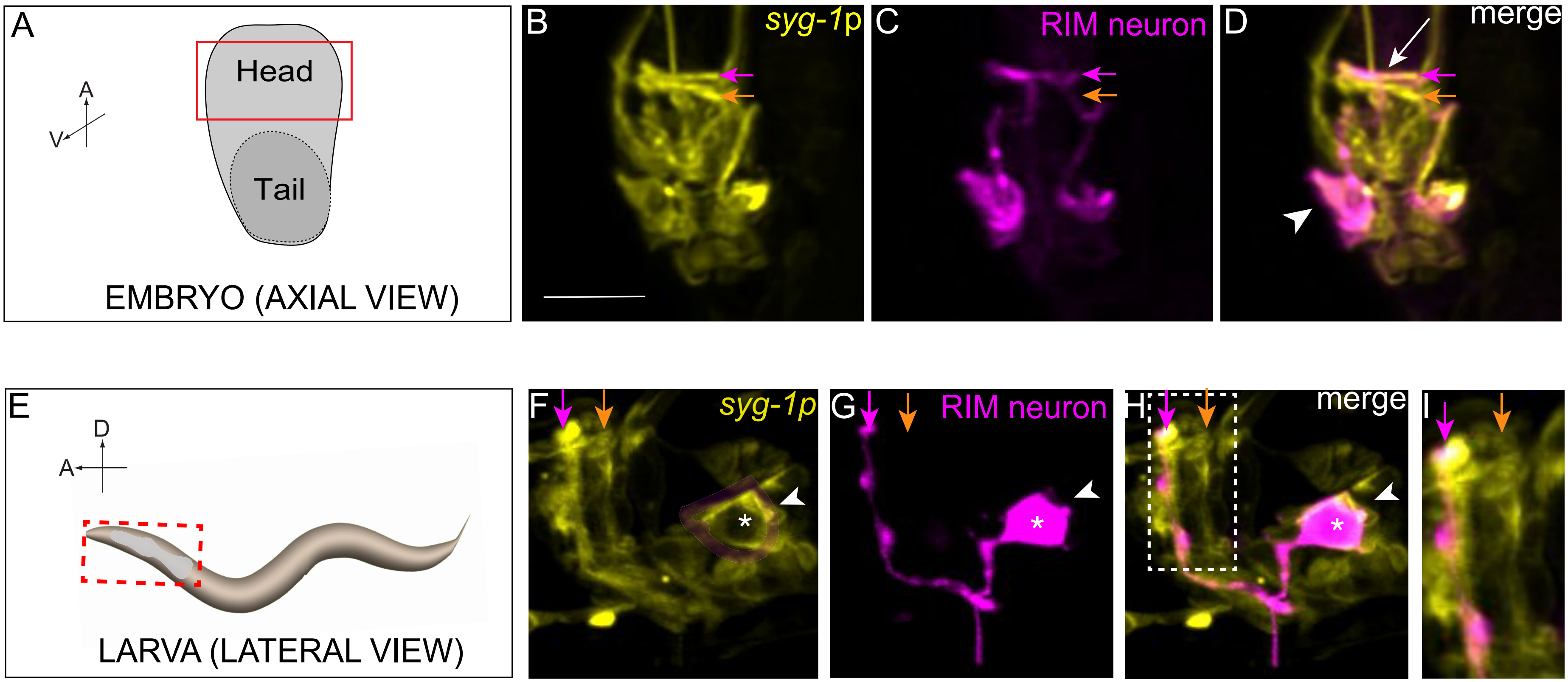

### Supplementary Fig. 7

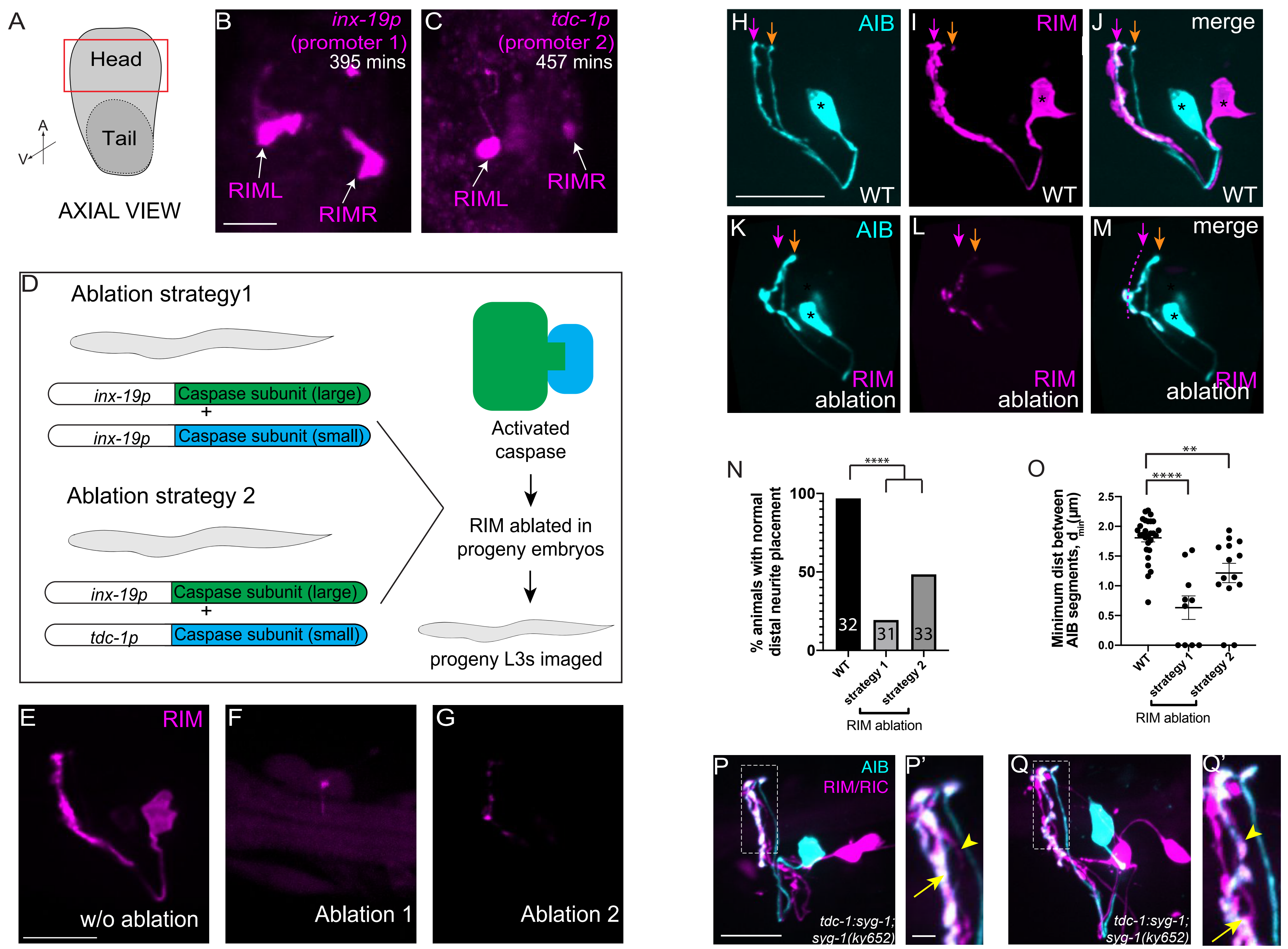

### Supplementary Fig. 8

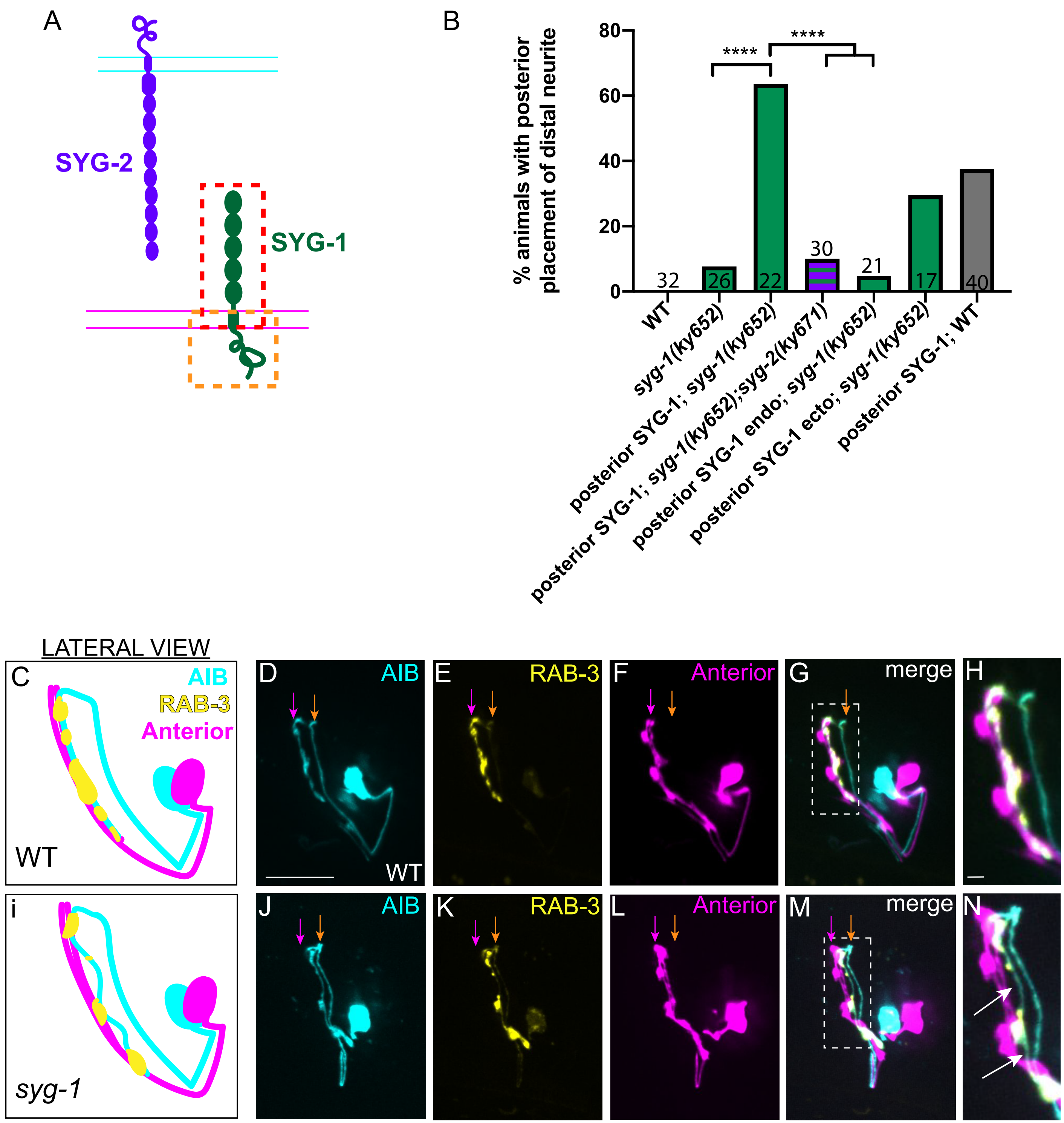

#### Supplementary Figure legends

##### Supplementary Fig. 1: Neurite positions of the bilaterally symmetric AIB neurons

**A-C,** Pseudo-colored confocal maximum intensity projections showing AIBL (**A**), AIBR (**B**) and merge (**C**). Orange and magenta arrows indicate positions of the proximal and distal neurites, positioned in the posterior and anterior neighborhoods respectively (see Fig. 1). Note that the proximal and distal neurites of AIBL and AIBR completely overlap in the lateral view, consistent with what would be expected based on their positions and projections from the EM reconstructions (in **D-I**; (White et al., 1986)). Scale bar = 10 μm applies **A-C**.

**D-F,** EM and *in vivo* fluorescent microscopy views of the AIBL-AIBR neuron pair. **(D)** Schematic of the AIBL-AIBR neuron pair in the context of the nerve ring (light neon); **(E)** Pseudo-colored confocal maximum intensity projections of AIBL (cyan) and AIBR (yellow) (3D projection of this dataset shown in Supplementary Movie 4); **(F)** Volumetric reconstructions of AIBL and AIBR from segmented EM datasets (JSH, (White et al., 1986)).

**G-I,** As **D-F**, but axial view.

In all images (**D-I**), arrowheads indicate the posterior-anterior shift of the two neurons crossing each other to form a chiasm. The grey circle in **G** depicts the pharynx. Scale bars = 10 μm in **E,H**, and 3 μm in **F,I**.

**Supplementary Fig. 2: AIB contacts, synaptic distribution and network properties enable its function as a connector hub neuron**

**A,** Volumetric reconstruction of AIBL (cyan), a posterior neighborhood neuron (ASEL, orange) and an anterior neighborhood neuron (RIMR, magenta) from an L1 (5 hours post hatching) and an adult connectome dataset (45 hours post hatching, (Witvliet et al., 2020)). Scale bars = 3 μm. Note that ASEL contacts AIBL exclusively in the posterior neighborhood, and RIMR contacts AIBL exclusively in the anterior neighborhood. AIB is similarly positioned into the same neighborhoods in all available connectome datasets examined (which span all larval developmental stages; data not shown; (White et al., 1986, Witvliet et al., 2020)). The observation that AIB is already positioned in the two neighborhoods in the L1 stage indicates that AIB placement occurs during embryogenesis.

**B,C,** Axial view of the AIB neurite and neuron-neuron contact areas between AIBL and anterior neighborhood neuron, RIMR (**B**) and AIBL and posterior neighborhood neuron, ASEL (**C**) from the segmented EM dataset of the L4 stage animal JSH (Brittin et al., 2018, White et al., 1986). Contacts are colored red (see Methods) and overlaid on 3D volumetric reconstructions of the AIB neurite (cyan). Orange and magenta arrows indicate the posterior and anterior neighborhoods respectively. Scale bar = 3 μm, applies to **B,C.**

**D,** Volumetric reconstruction of AIBR from the JSH electron microscopy connectome dataset (White et al., 1986) in lateral view. Postsynaptic (red) and presynaptic (yellow) regions of the neurite, based on synaptic connectivity maps of AIBR, are indicated. Note that the postsynaptic and presynaptic regions coincide with the proximal and distal segment of AIB. Arrowhead points to the chiasm. Scale bar = 1 μm

**E-G,** Representative confocal image showing a lateral view of an AIB neuron with postsynaptic sites (red, labeled by GLR-1:GFP, **E**) and presynaptic sites (yellow, labeled by mCh:RAB-3, **F**). **G** is a merge of **E** and **F**. Note the opposite polarity in the posterior and anterior neighborhoods (indicated by the orange and magenta arrows respectively)

Scale bar = 10 μm, applies to **E-G**.

**H,** From the available segmented serial section EM and neuron-neuron contact data (Brittin et al., 2021, Moyle et al., 2021, White et al., 1986, Witvliet et al., 2020), neuron-neuron adjacency matrices were generated as depicted in the schematic.

**I,** Cosine similarity plot for AIBR. The cosine similarity values (Han et al., 2012) of AIB contacts between each pair of connectome datasets (White et al., 1986, Witvliet et al., 2020) are plotted as a heat map with the color bar indicating the values corresponding to the shades. The similarity values are >0.5 for all pairs of connectome datasets, indicating positive correlation between distribution of AIB contacts in different datasets. This suggests that distribution of AIB contacts with other neurons is largely established in early larval stages (L1) and maintained through development. The labels assigned to the connectome datasets comprise of the developmental stage and name of the animal sectioned (all times are measured post hatching) (White et al., 1986, Witvliet et al., 2020).

**J,** Box plot (10 to 90 percentile) of betweenness centrality values for an L1 (5 hr post hatching) connectome dataset and an adult connectome dataset (45 hr post hatching,

(Witvliet et al., 2020)). The gray dots represent neurons whose centrality values lie above the 90-percentile mark or below the 10-percentile mark. The betweenness centrality values (see Methods) for AIBL and AIBR (indicated by cyan and yellow stars respectively) lie above the 90-percentile mark in both datasets. High betweenness centrality being a standard property of rich-club neurons (Towlson et al., 2013), this indicates that contacts of AIBL and AIBR exhibit rich-club features from early developmental stages (L1) to adulthood.

**Supplementary Fig. 3: Strategies for labeling and visualizing AIB outgrowth dynamics in early embryos**

**A,** Schematic of the ZIF-1/1-ZF1 degradation system used for subtractive fluorescent labeling of specific neurons in embryos (Armenti et al., 2014), also see Methods. Briefly we used *lim-4p*, expressed in sublateral neurons (Santella et al., 2015) to express ZIF-1 and *unc-42p*, expressed in the sublateral neurons + AIB + ASH (http://promoters.wormguides.org) to express ZF1-tagged PH:GFP. This results in degradation of PH:GFP from the sublateral neurons, resulting in cell-specific labeling of AIB and/or the ASH neurons.

**B,** diSPIM image showing *unc-42p*-driven membrane-tethered PH:GFP expression, without ZIF-1/ZF1 mediated degradation, in neurons of the embryonic nerve ring. Distinction of AIB neurite outgrowth dynamics is not possible in this background due to abundant labeling. Scale bar = 10 μm applies to **B,C.**

**C,** diSPIM image showing the nerve ring of an embryo expressing the ZIF-1/ZF1 degradation construct strategy outlined in **A**. The identity of AIB was confirmed by colocalization and lineaging as described (Moyle et al., 2021).

**D,E,** Imaging methods that we established and implemented for the investigation of neurodevelopmental events in *C. elegans* embryos. **D,** A triple-view line-scanning confocal microscope that provides enhanced (2-fold) axial resolution compared to conventional confocal microscopy (Wu et al., in prep). To image AIB in living nematode embryos, we additionally created a two-step deep learning framework that denoises the raw data, enabling us to turn down the illumination intensity ~30-fold, offering more gentle imaging than conventional confocal microscopy. **E,** Dual-view inverted selective plane illumination microscopy (diSPIM), a light-sheet microscopy technique for long-term imaging of AIB neurite development. Deconvolution and fusion of images from orthogonal views result in isotropic spatial resolution.

**F-H**, Time-lapse showing relative positions of AIBL (pseudo-colored in cyan) and AIBR (pseudocolored in yellow) in the embryonic nerve ring. Images are reconstructions derived from triple-view line-scanning confocal microscopy, which used a deep learning algorithm for denoising and deconvolving all 3 views. The dashed white lines represent the dorsal midline of the nerve ring. The dotted boxes represent the dorsal half of the nerve ring and are magnified in **F'-H'**. **F''-H''** are schematic diagrams representing the images in **F-H**. In (**F,F'F''**), the neurites are initially positioned in the same neighborhood. In (**G,G',G''**) they have separated partially from the tip up to a point along their lengths (arrow). In (**H,H',H''**) they have separated completely up to the dorsal midline (arrowhead). The high spatial resolution allows us to clearly distinguish the two neurites and determine their relative positions reliably, confirming results in Fig. 2 and enabling detailed quantifications. Scale bar = 2 μm in **F**, applies to **G,H,** and 1 μm in **F'**, applies to **G',H'**.

**I,** Schematic showing AIBL and AIBR in the context of the nerve ring (light neon), pharynx (grey) and the anterior and posterior neighborhood (magenta and orange regions). AIBL exits the posterior neighborhood (direction of outgrowth indicated by black arrow) and cuts through the nerve ring to meet the anterior neighborhood. $\alpha$ is the angle of exit. AIBR also exits similarly. We also measured $\beta$: the angle between tangents drawn at the point of downward bend of the nerve ring in the posterior neighborhood, as indicated in schematic (from embryos in which posterior neighborhood neurons are labeled by *nphp-4p*:PH:GFP).

**J,** Scatter plot of $\alpha$ and $\beta$ values (n=6, 3 AIBL and 3 AIBR neurons measured from 3 embryos for each of $\alpha$ and $\beta$). Unpaired two-tailed t test indicates no significant difference (n.s.) between $\alpha$ and $\beta$ values (p=0.1368). The AIB distal neurite therefore exits tangentially from the posterior neighborhood, consistent with AIB losing adhesion in this neighborhood, growing straight instead of following the arc of the nerve ring and crossing the nerve ring towards its eventual encounter with the RIM neuron in the anterior neighborhood.

**Supplementary Fig. 4: SYG-1 and SYG-2 regulate placement of the AIB neurite specifically in the anterior neighborhood**

**A,** Scatter plot of lengths of the dorsal midline shift (that form a chiasm for the neuron pair, see Fig. 1 and Supplementary Fig. 1) for wild type (n=18) and *syg-1 (ky652)* (n=18). Error bars indicate standard error of the mean (S.E.M.). ***p=0.0008 (unpaired two-tailed t-test). In wildtype animals, the chiasm is stereotyped and similar in length across L4 stage animals, as measured from confocal micrographs and displayed in this scatter plot (mean length = 2.97±0.05 μm, n =18), and electron micrographs (dorsal midline shift length in AIBL and AIBR in electron micrographs of an L4 stage animal, JSH, are 3.01 μm and 3.16 μm respectively). In *syg-1(ky652)*, the mean length of the chiasm is significantly smaller and is = 1.96±0.27 μm (n=18). n represents the number of AIB neurons measured, applies to **B,C.** Effect size estimate, d = 1.233.

**B,** Scatter plot of distal neurite lengths for wild type (n=31) and *syg-1(ky652)* (n=30).

For quantifications in **A** and **B**, same images as for Fig. 4M were used.

**C,** Scatter plot of nerve ring width as measured from strains expressing a nerve ring marker *cnd-1p:PH:GFP* (see Methods), in WT (n=14) and *syg-1(ky652)* (n=14) backgrounds. Two values of nerve ring width were obtained from each animal (one from each side). n = number of animals of each genotype from which measurements were done.

For **B** and **C**, unpaired two-tailed t test indicates no significant (abbreviated by n.s.) difference (p = 0.0793 and 0.3140 respectively). Error bars indicate standard error of the mean (S.E.M.).

**D,E,** Representative confocal images of a *syg-1(ky652)* (**D**) and a *syg-2(ky671)* (**E**) animal with AIB labeled with cytoplasmic mCherry (cyan) and the posterior neighborhood markers, the AWC and ASE neurons labeled with cytoplasmic GFP (orange). Note that the placement of the AIB neurite in the posterior neighborhood is unaffected in *syg-1(ky652)* and *syg-2(ky671)*. The orange and magenta arrows indicate the positions of the posterior and anterior neighborhoods respectively. Scale bar = 10 μm.

**F-H'** Representative confocal images of AIB (cyan) and the anterior neighborhood (magenta) in strains with cell-specific SYG-2 expression (**F**), cell-specific SYG-1 expression (**G**) and a cosmid containing *syg-1* (recapitulating endogenous SYG-1 expression) (**H**). The dashed boxes in **F**,**G** and **H** represent the region of contact between AIB and the anterior neighborhood, magnified in **F'**, **G'** and **H'** respectively. Note complete alignment of the AIB distal neurite with the anterior neighborhood in **F'** and **H'** and detachment in **G'**.

Scale bar = 10 μm for **F**,**G**,**H**, and 1 μm for **F'**,**G'**,**H'**.

**Supplementary Fig. 5: Spatiotemporal regulation of *syg-1* transcriptional reporter expression during embryogenesis**

**A-D,** Time-lapse images of *syg-1* reporter expression during embryogenesis (450-630 m.p.f.). Images are deconvolved diSPIM maximum intensity projections. The dashed boxes represent the dorsal half of the nerve ring and are magnified in **E-H**. **E'-H'** are schematic diagrams representing the images in **E-H**. In (**A,E,E'**), *syg-1* expression is primarily visible in a single band containing amphid neurites, and therefore coincident with the AIB posterior neighborhood (indicated with orange arrow). The magenta dashed line and magenta arrows point to the anterior neighborhood. (**B,F,F'**) show onset of weak *syg-1* expression in the anterior neighborhood (white arrow in **F**) and ingrowth of *syg-1*-expressing neurites into this neighborhood (white arrowhead, identified as RIM neurons by colocalization, see Supplementary Fig. 6). *syg-1* expression increases in the anterior neighborhood and decreases in the posterior neighborhood as embryonic development progresses (**C**,**G**,**G'**,**D**,**H**,**H'**), quantified in Fig. 5S and similar to the SYG-1 protein reporter (Fig. 5J-R').

Scale bar = 10 μm in **A-D**.

Scale bar = 1 μm in **E-H**.

All times are in m.p.f. (minutes post fertilization).

**Supplementary Fig. 6: SYG-1 is expressed in the anterior neighborhood RIM neurons**

**A,** Schematic of the axial view of an embryo with the red box showing the region in the head containing the nerve ring.

**B-D,** Deconvolved diSPIM maximum intensity image of (**B**) membrane-targeted PH:GFP driven by the *syg-1* promoter (as in Supplementary Fig. 5) and (**C**) RIM in an embryo. **d** is a merge of **B** and **C**. Note the colocalization of the RIM neurites (arrow) and the RIM cell body (arrowhead) in **D** with the *syg-1* reporter in the anterior neighborhood.

**E-I,** Expression of the same *syg-1* reporter as in **B-D,** but in larval stage 3 in a lateral view (**E**). The *syg-1* reporter (**F**) is co-expressed with a cytoplasmic RIM neuron marker (**G**). **H** is a merge of **F** and **G**. The dashed box represents the region of the nerve ring containing the RIM neuron and the *syg-1*-expressing neurons. Note the RIM neurite colocalizes with the anterior band of *syg-1* expression, coincident with the AIB anterior neighborhood (magenta arrow). The white arrowhead in **F-H** and semi-transparent magenta outline in **F** indicates colocalization of the RIM cell body with the *syg-1* reporter.

Scale bar = 10 μm applies to **B-I**.

**Supplementary Fig. 7: The RIM neurons regulate AIB distal neurite placement**

**A,** Schematic of the axial view of an embryo where the red box highlights the region in the head where nerve ring neurons are present, and cropped from images of whole embryos, to produce the images in **B,C.**

**B,C,** diSPIM maximum intensity projections of fluorescently labeled RIM neurons (arrows) in embryos prior to AIB distal neurite placement (~500-550 m.p.f., minutes post fertilization), labeled with *inx-19*p:GFP (**B**) or *tdc-1*p:GFP (**C**). Scale bar = 10 μm applies to **B,C**.

**D,** Schematic showing strategies used for ablation of the RIM neurons in embryos. In strategy 1, a small (p12) and a large (p17) subunit of human Caspase-3 are both expressed by *inx-19*p, similar to previously described (Chelur and Chalfie, 2007). The *inx-19*p is expressed in the RIM neurons from 370 m.p.f – the time of their birth. In strategy 2, p12 is expressed by *inx-19*p and p17 by *tdc-1*p (*tdc-1*p is expressed in the RIM neurons ~445 m.p.f.). These caspase subunits are therefore expected to reconstitute expression (and induce ablation) in embryonic RIM neurons.

**E,** Representative confocal image of a wild type L3 animal expressing membrane-targeted PH:GFP in the RIM neurons with RIM-specific promoter *gcy-13*p.

**F,G** As **E**, but in animals additionally expressing the caspase subunits for (**f**) ablation strategy 1 and (**G**) ablation strategy 2. Note the absence of RIM labeling, indicating successful ablation of the RIM neurons.

Scale bar = 10 μm, applies to **E-G**.

**H-M,** Confocal images showing AIB (labeled with cytoplasmic mCherry; **H,K**) and RIM neurons (labeled with PH:GFP; **I,L**) and merged images (**J,M**) for wild type animals (**H-J**) and animals in which RIM was genetically ablated (**K-M**). RIM ablation was achieved using Strategy 2, see Methods. Magenta dashed line (**M**) represents AIB anterior neighborhood.

**N,** Quantification of the penetrance of the AIB neurite placement defect as the percentage of animals with normal AIB neurite placement in the anterior neighborhood. Strategy 1 and Strategy 2 refer to split caspase ablations (Chelur and Chalfie, 2007) using two different combinations of promoters expressed in RIM neurons (see Methods). ****$p<0.0001$ (two-sided Fisher's exact test). Numbers on bars represent number of animals examined.

**O,** Quantification of the minimum perpendicular distances between the AIB proximal and distal neurites in WT (n=28) and RIM-ablated populations (n=10 for strategy 1 and n=14 for strategy 2). ****$p<0.0001$; **$p=0.0011$ (one-way ANOVA with Dunnett's multiple comparisons test). n represents the number of AIB neurons measured from 14, 5 and 7 animals from the WT, ablation strategy 1 and ablation strategy 2 populations respectively. Effect size estimate, d = 2.313.

**P,Q,** Confocal micrographs of animals where AIB (cyan) and the RIM and RIC neurons (magenta) are co-labeled and SYG-1 is expressed specifically in RIM and RIC (see Methods) in a *syg-1(ky652)* mutant background. The dashed box represents the nerve ring region containing the neurites of AIB, RIM and RIC, and is magnified in **P'** and **Q'**. The AIB distal neurite is positioned along RIM (**P'**) or along both RIM and RIC (**Q'**). The yellow arrowheads and yellow arrows point at the RIC neurite and the RIM neurite respectively.

Scale bar = 10 μm for **H-M**, **P,Q,** and 2 μm for **P'** and **Q'**.

**Supplementary Fig. 8: An extracellular SYG-1-SYG-2 interaction regulates AIB neurite placement and presynaptic localization**

**A,** Schematic of the receptor-ligand pair SYG-1 (green) and SYG-2 (purple). The red dashed box includes the SYG-1 extracellular Ig domains and transmembrane domain (collectively referred to as SYG-1 ecto). The yellow dashed box includes the SYG-1 transmembrane domain and cytoplasmic domains (collectively referred to as the SYG-1 endodomain or SYG-1 endo).

**B,** Quantification of penetrance of the ectopic AIB neurite placement as the percentage of animals with the AIB distal neurite partially positioned in the posterior neighborhood in the indicated genotypes. ****p<0.0001 (by two-sided Fisher's exact test) for *syg-1(ky652)* and *mgl-1bp* expressed SYG-1 (SYG1 expression specifically in the posterior neighborhood, denoted as posterior SYG-1), for posterior SYG-1 expression in *syg-1(ky652)* background and *syg-1(ky652);syg-2(ky671)* background, and for posterior

SYG-1 expression and posterior SYG-1 endodomain (only) expression in *syg-1(ky652)* background. Number on bars represent the number of animals examined. The first three bars are the same as the ones corresponding to these genotypes in Fig. 6K.

**C-H,** Schematic (**C**) and representative confocal image of the AIB neurite (**D**), AIB presynaptic sites (**E**) and neurite of the postsynaptic partner, the RIM neuron in the anterior neighborhood (**F**). **G** is a merged image. The dashed box represents the region of contact between the AIB and RIM neurites, magnified in **H**.

**I-N,** As **C-H** but in the *syg-1(ky652)* mutant background. Note the gaps between the AIB distal neurite and the RIM neurites (**N**) and reduced localization of RAB-3 along the AIB neurite in the region where it is detached from RIM (white arrows in **N**).

Scale bar = 10 μm to **D-H**, **J-N**. The black, green and purple bars represent WT, *syg-1(ky652)* and *syg-2(ky671)* backgrounds, respectively.

#### Supplementary Tables

**Supplementary Table 1. Single-cell RNAseq dynamics are consistent with our observed expression changes,** related to Fig. 5 and Supplementary Fig. 5

SYG-1 single-cell RNAseq values (Packer et al., 2019) for neurons belonging to AIB's posterior and anterior neighborhoods, at time points capturing the AIB distal neurite transition (430-550 and 550-690 m.p.f). Values are adjusted transcripts per million (t.p.m) estimates where ci.95p >0 (Packer et al., 2019). Green triangles indicate an increase in expression over time, red triangles indicate a decrease. For each neighborhood, neurons are listed in descending order of the highest number of contacts with AIBL and AIBR in JSH and N2U EM datasets (White et al., 1986). Neurons with >20 documented contacts in JSH and N2U EM reconstructions were included.

| Neuron Class | Adjusted tpm estimate for *syg-1* transcripts (Packer et al). | | |
|---|---|---|---|
| | (a) 430-550 m.p.f | (b) 550—690 m.p.f | Change in expression (b-a) |
| | AIB posterior neighborhood | | |
| ADL | 545.3 | 13.1 | -532.2 |
| ASH | 105.9 | 61.9 | -44 |
| ASK | 39.5 | 7.8 | -31.7 |
| AFD | 144.3 | 229.4 | 85.1 |
| AIY | 1491.9 | 180.3 | -1311.6 |
| | AIB anterior neighborhood | | |
| RIM | 1536.3 | 1078.2 | -458.1 |
| AVE | 333.8 | 1246.5 | 912.7 |
| SMD | 837.4 | 168.9 | -668.5 |
| AVK | 7.3 | 34 | 26.7 |
| AVB | 0 | 419.2 | 419.2 |
| SIA | 199.3 | 304.4 | 105.1 |
| RIV | 588.8 | 1367.1 | 778.3 |

**Supplementary Table 2. Strains generated and used in this study**

| **Strain identifier** | **Source** | **Genotype** |
|---|---|---|
| BV276 | (Duncan et al., 2019) | ujIs113[pie-1p::mCherry::H2B::pie-1 3'UTR + nhr-2p::his-24::mCherry::let-858 3'UTR + unc-119(+)];<br>II |
| JIM158 | gift from John Murray | ujIs113;oyIs48[Pceh-36::GFP, lin-15(+)];V |
| DCR5516 | This paper | olaIs67[DACR2245 at 40 ng/uL+DACR1412 at 30 ng/uL+DACR218 at 30 ng/uL];X |
| DCR5761 | This paper | olaex3394[DACR2233 at 60 ng/uL+DACR2651 at 60 ng/uL+DACR218 at 30 ng/uL] |
| DCR6222 | This paper | olaex3666[DACR199 at 2 ng/uL+ DACR218 at 30 ng/uL];olaIs67 |
| DCR6301 | This paper | oyIs48;ujIs113;olaIs67 |
| DCR7648 | This paper | olaex4624[DACR3149 at 10 ng/uL+DACR218 at 30 ng/uL];olaIs67 |
| DCR5517 | This paper | olaIs68[DACR2245 at 40 ng/uL+DACR1412 at 30 ng/uL+DACR218 at 30 ng/uL] |
| DCR8220 | This paper | olaIs68;*syg-1(ky652)* |
| DCR8486 | This paper | olaIs68;*syg-1(ok3640)* |
| DCR8183 | This paper | olaex4624;olaIs68;*syg-1(ky652)* |
| CX5862 | This paper | kyIs235;kyEx679;*syg-1(ky652)* |
| DCR8180 | This paper | kyEx679;olaIs68;*syg-1(ky652)* |
| DCR8489 | This paper | olaex4624;kyEx679;olaIs68;*syg-1(ky652)* |
| DCR6767 | This paper | olaIs68;*syg-2(ky671)* |
| DCR8468 | This paper | olaex4624;olaIs68;*syg-2(ky671)* |
| DCR8488 | This paper | oyIs48; olaIs68;*syg-1(ky652)* |
| DCR8440 | This paper | olaex5120[DACR3529 at 30 ng/uL+DACR1412 at 30 ng/uL+DACR218 at 30 ng/uL] |
| DCR8365 | This paper | olaex5063[DACR3492 at 25 ng/uL+DACR3505 at 40 ng/uL+DACR2312 at 25 ng/uL+DACR20 at 25 ng/uL];olaIs67 |
| DCR6814 | This paper | olaex4071[DACR2637 at 15 ng/uL+DACR218 at 30 ng/uL];olaIs67 |
| DCR6920 | This paper | olaex4130[DACR2704 at 100 ng/uL+DACR218 at 50 ng/uL];ujIs113 |
| DCR6782 | This paper | olaex4052[DACR2607 at 100 ng/uL+DACR2609 at 25 ng/uL+DACR218 at 30 ng/uL];olaIs67 |
| DCR6784 | This paper | olaex4054[DACR2607 at 100 ng/uL+DACR2609 at 25 ng/uL+DACR218 at 30 ng/uL];olaIs67 |
| DCR5730 | This paper | olaex3388[DACR2371 at 75 ng/uL+DACR2404 at 30 ng/uL+DACR218 at 30 ng/uL] |

| | | |
|---|---|---|
| DCR7642 | This paper | olaex4618[DACR2607 at 100 ng/uL+DACR2609 at 25 ng/uL+DACR2863 at 25 ng/uL+DACR218 at 30 ng/uL];olaIs67 |
| DCR7643 | This paper | olaex4619[DACR2607 at 100 ng/uL+DACR2609 at 25 ng/uL+DACR2863 at 25 ng/uL+DACR218 at 30 ng/uL];olaIs67 |
| DCR6633 | (Moyle et al., 2021) | olaex3949[DACR2607 at 100 ng/uL+DACR2609 at 25 ng/uL+DACR2351 at 25 ng/uL+DACR218 at 30 ng/uL] |
| DCR4894 | This paper | olaex2887[DACR2245 at 100 ng/uL+DACR2404 at 30 ng/uL+DACR218 at 30 ng/uL] |
| DCR6082 | This paper | olaex3570[DACR2481 at 10 ng/uL+DACR218 at 50 ng/uL];ujIs113 |
| DCR8421 | This paper | olaex5105[DACR3605 at 50 ng/uL+ DACR218 at 30 ng/uL] |
| DCR8347 | This paper | olaIs117[DACR3502 at 30 ng/uL+DACR20 at 25 ng/uL];olaIs68 |
| DCR8350 | This paper | olaIs117;olaIs68;*syg-1(ky652)* |
| DCR8361 | This paper | olaex5059[DACR3503 at 10 ng/uL+DACR20 at 25 ng/uL];olaIs68;*syg-1(ky652)* |
| DCR8352 | This paper | olaex5050[DACR3698 at 30 ng/uL+DACR20 at 25 ng/uL];olaIs68;*syg-1(ky652)* |
| DCR8470 | This paper | oyIs48; olaex5059; olaIs68; *syg-1(ky652)* |
| DCR8472 | This paper | oyIs48; olaIs117; olaIs68; *syg-1(ky652)* |
| DCR6841 | This paper | olaex4087[DACR1412 at 30 ng/uL+DACR2618 at 50 ng/uL+DACR218 at 30 ng/uL] |
| DCR8758 | This paper | ujIs113;oyIs48;olaIs68;*syg-2(ky671);* |
| DCR8762 | This paper | olaEx5279[DACR3527 at 30 ng/uL+DACR20 at 25 ng/uL]; olaIs68; *syg-1(ky652)* |
| DCR8759 | This paper | olaEx5276 [DACR3780 at 5 ng/ul+DACR1412 at 20 ng/uL+DACR218 at 30 ng/uL] |
| DCR8764 | This paper | olaEx5281[Pinx-1::syg-2b::unc-54UTR at 30 ng/uL+DACR20 at 30 ng/uL]; olaIs68; *syg-2(ky671)* |
| DCR8766 | This paper | olaEx5283[DACR3781 at 30 ng/uL+DACR20 at 25 ng/uL]; olaIs68;*syg-1(ky652)* |
| DCR8767 | This paper | olaIs117; olaIs68; *syg-1(ky652); syg-2(ky671)* |
| BV293 | (Fan et al., 2019) | zbIs3[cnd-1p::PH::GFP] |
| DCR8772 | This paper | zbIs3;olaIs68;*syg-1(ky652)* |

**Supplementary Table 3. Plasmids generated and used in this study**

| Plasmid name | Source | Details |
|---|---|---|
| DACR1412 | (Moyle et al., 2020) | Pinx-1::mCherry::unc-54UTR |
| DACR2245 | (Moyle et al., 2020) | Pinx-1::eGFP::rab-3::unc-54UTR |
| DACR2233 | This paper | Pinx-1::glr-1::GFP::unc-54UTR |
| DACR2651 | This paper | Pinx-1::mCh::rab-3::unc-54UTR |
| DACR199 | This paper | Pcex-1::GFP |
| DACR3149 | This paper | Pcex-1::mTagBFP1::unc-54UTR |
| DACR3529 | This paper | Psyg-1::PHD::GFP::unc-54UTR |
| DACR3492 | This paper | Pinx-19::p12-caspase3::unc-54UTR |
| DACR3493 | This paper | Pinx-19::p17-caspase3::unc-54UTR |
| DACR3505 | This paper | Ptdc-1::p17-caspase3::unc-54UTR |
| DACR2637 | This paper | Ptdc-1::GFP::unc-54UTR |
| DACR2863 | This paper | Ptdc-1::PHD::GFP::unc-54UTR |
| DACR2704 | This paper | Pinx-19::PHD::GFP::unc-54UTR |
| DACR2481 | This paper | Pnphp-4::PHD::GFP::unc-54UTR |
| DACR2404 | This paper | Pinx-1::mCh::PHD::unc-54UTR |
| DACR2371 | This paper | Punc-42::PHD::GFP::unc-54UTR |
| DACR2607 | (Moyle et al., 2020) | Punc-42::ZF1::PHD::GFP::unc-54UTR |
| DACR2609 | (Moyle et al., 2020) | Plim-4(includes exons 1-3)::SL2::Zif-1::unc-54UTR |
| DACR2351 | (Moyle et al., 2020) | Plim-4(includes exons 1-3)::mCherry::unc-54UTR |
| DACR3605 | This paper | Ptdc-1::mScarlet::PHD::unc-54UTR |
| DACR3502 | This paper | Pmgl-1b::syg-1b::unc-54UTR |
| DACR3503 | This paper | Pnphp-4::syg-1b::unc-54UTR |
| DACR3698 | This paper | Pmgl-1b::syg1ecto::unc-54UTR |
| DACR2618 | This paper | Pinx-1::cla-1::GFP::unc-54UTR |
| DACR3527 | This paper | Pinx-1::syg-1b::unc-54UTR |
| DACR3780 | This paper | Psyg-1::syg-1::GFP::unc-54UTR |
| DACR3781 | This paper | Pmgl-1b::syg-1endo::unc-54UTR |

#### Supplementary Movies

**Supplementary Movie 1. Neighborhoods of AIB**, related to Fig. 1, Supplementary Fig. 1 and 2

3D volumetric reconstruction of AIBR (cyan), posterior neighborhood neuron AWCR (orange) and anterior neighborhood neuron, RIML (magenta) from segmented EM micrographs from the JSH EM dataset (White et al., 1986). Scale bar = 2 μm.

**Supplementary Movie 2. AIB and anterior neighborhood neuron, RIM,** related to Fig. 1 and Supplementary Fig. 2

3D projection confocal image of AIB (cyan) and the RIM neurons (magenta) of the anterior neighborhood. The RIM neurites show extensive overlap with the AIB distal neurite. This movie was made from the same image as in Fig. 1D,I. Scale bar = 10 μm.

**Supplementary Movie 3. AIB and posterior neighborhood neurons,** related to Fig. 1 and Supplementary Fig. 2

3D projection confocal image of AIB (cyan) and posterior neighborhood amphid sensory neurons (orange). The posterior neighborhood neurons show extensive overlap with the AIB proximal neurite. This movie was made from the same image as in Fig. 1E,J. Scale bar = 10 μm.

**Supplementary Movie 4. The bilaterally symmetric AIBL and AIBR neurons**, related to Fig. 1 and Supplementary Fig. 1

3D projection confocal image of the AIB neuron pair, AIBL (cyan) and AIBR (yellow), pseudocolored to distinguish the AIBL and AIBR neurites and their chiasm at the dorsal midline (in the video, the crossover at the very top). The maximum intensity projections in Supplementary Fig. 1E,H are from this same dataset. Scale bar = 10 μm.

**Supplementary Movie 5. Polarized localization of AIB presynaptic sites**, related to Supplementary Fig. 2

3D projection confocal image of AIBR (cyan) with presynaptic sites labeled with a CLA-1 reporter (yellow) (Xuan et al., 2017). Posterior and anterior neighborhoods are marked in the lateral and axial views with orange and magenta arrows, respectively. Note localization of CLA-1 (yellow) specifically along the distal neurite. Scale bar = 10 μm

**Supplementary Movie 6. Selective AIB labeling by Zif-1/ZF1 mediated degradation**, related to Fig. 2 and Supplementary Fig. 3

diSPIM maximum intensity projection images of embryos, labeled with membrane-tagged GFP (expression driven by an *unc-42* promoter, see Methods, (Armenti et al., 2014)) without (left) and with (right) Zif-1-ZF1 mediated degradation. The dataset with the Zif-1-ZF1 degradation was used in Supplementary Fig. 3C. Scale bar = 10 μm.

**Supplementary Movie 7. Layered expression of SYG-1 in the nerve ring in embryos**, related to Fig. 5 and Supplementary Fig. 5

3D projection of deconvolved diSPIM images of an embryo expressing the *syg-1* transcriptional reporter (*syg-1p*) (this movie is the same as the 529 m.p.f. timepoint in

Supplementary Fig. 5, also see Methods). Expression of *syg-1p* is restricted to two nerve ring neighborhoods, corresponding to the AIB posterior and anterior neighborhoods (as indicated in Fig. 5). Scale bar = 10 μm.

**Supplementary Note – Biophysical equations for retrograde zippering and unzippering**

Force balance equations to derive velocities at the zippering and unzippering points

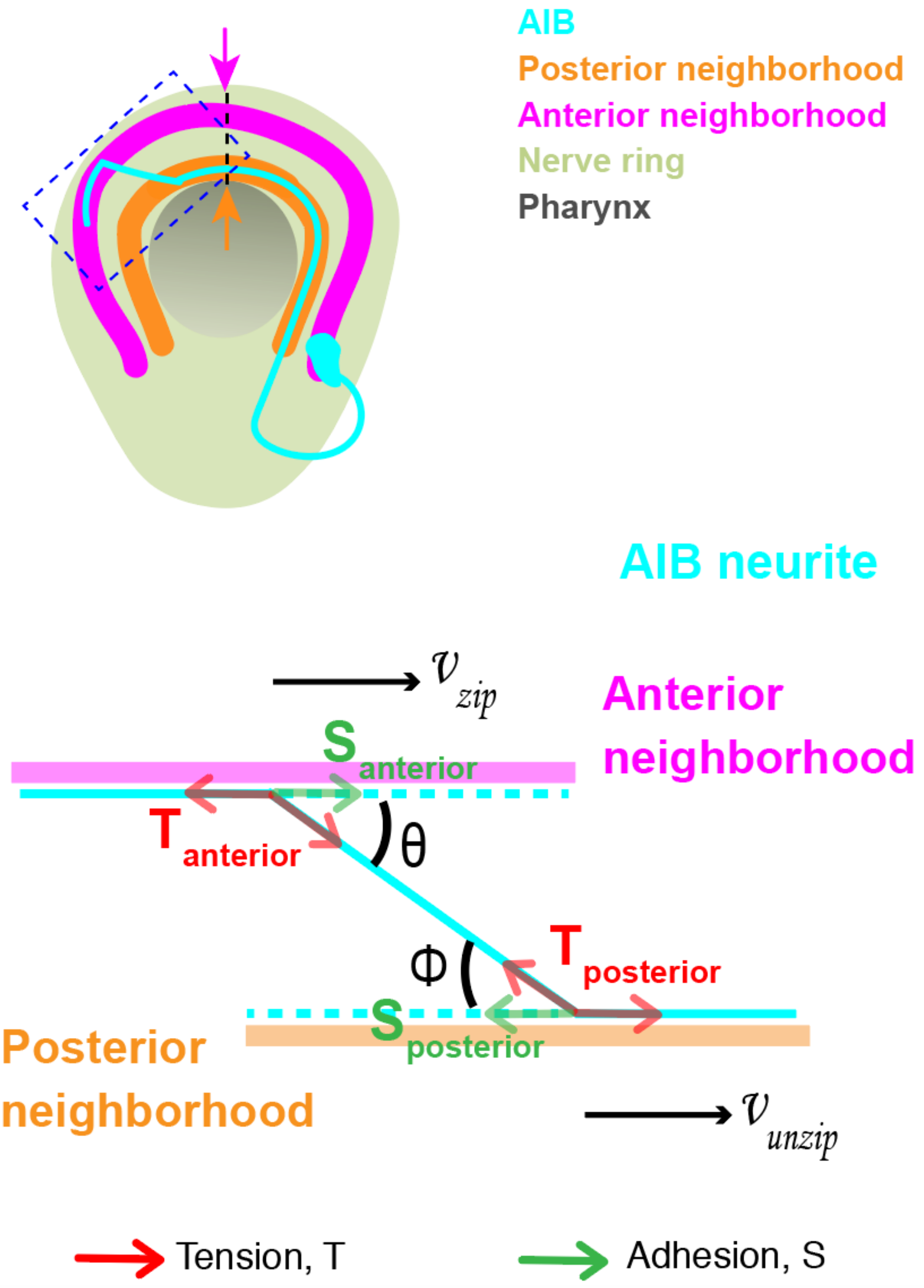

According to the biophysical models proposed for tissue culture cells (Smit et al., 2017), zippering and unzippering of neurite shafts occur as a result of primarily two forces acting on the neurite: adhesion, S and tension, T. Additionally as the neurite zippers or unzippers, it experiences frictional forces that act opposite to the direction of movement of the neurite, i.e., opposite to the velocity of the neurite.

At the zippering fork (see schematic above), adhesion acts in the direction of zippering, i.e., (in the direction of the zippering velocity $v_{zip}$) and favors zippering, while tension and friction act in the opposite direction, disfavoring zippering. Assuming dynamic equilibrium (Smit et al., 2017), all the forces (with the + or – signs representing their directions) add up to 0, resulting in the following equation:

**Equation 1:**

$$S_{anterior} + T_{anterior} cos\theta - T_{anterior} - \eta v_{zip} = 0$$

where $v_{zip}$ = velocity of retrograde zippering, $S_{anterior}$ = adhesion of the AIB neurite to the anterior neighborhood, $T_{anterior}$ = mechanical tension along the neurite, $\eta v_{zip}$ = friction forces at the zipper fork and $\theta$ = zipper angle.

Therefore, zippering velocity, $v_{zip}$ would be given by:

**Equation 2:**

$$v_{zip} = \frac{S_{anterior}}{\eta} - \frac{T_{anterior}}{\eta}(1 - cos\theta)$$

On the other hand, at the unzippering fork (see schematic above), adhesion and friction act in the direction opposite to unzippering (i.e., opposite to the unzippering velocity $v_{unzip}$) and disfavors unzippering, while tension acts in the same direction favoring unzippering. Again, assuming dynamic equilibrium (Smit et al., 2017), all the forces (with the + or – signs representing their directions) add up to 0, resulting in the following equation: (Smit et al., 2017)

**Equation 3:**

$$S_{posterior} + T_{posterior} cos\varphi - T_{posterior} + \eta v_{unzip} = 0$$

Therefore, the velocity of unzippering from the posterior neighborhood, $v_{unzip}$, would be given by:

**Equation 4:**

$$v_{unzip} = -\frac{S_{posterior}}{\eta} + \frac{T_{posterior}}{\eta}(1 - cos\varphi)$$

where $v_{unzip}$ = velocity of unzippering, $S_{posterior}$ = adhesion of the AIB neurite to the posterior neighborhood, $T_{posterior}$ = mechanical tension along the neurite, $\eta v_{unzip}$ = friction forces at the unzippering fork and $\varphi$ = zipper angle.

Since the nerve bundles constituting the posterior and anterior neighborhoods are parallel near the dorsal midline, therefore,

**Equation 5:**

$$\theta = \varphi$$

Combining equations (2), (4) and (5), we get,

**Equation 6:**

$$v_{zip} + v_{unzip} = \frac{(S_{anterior} - S_{posterior})}{\eta} - \frac{(T_{anterior} - T_{posterior})}{\eta}(1 - cos\theta)$$

Since the same stretch of the AIB neurite that zippers onto the anterior neighborhood, concurrently unzippers from the posterior neighborhood (Fig 3C), and assuming mechanical tension is uniformly redistributed along the neurite (Smit et al., 2017), tension at the zippering and unzippering forks would be equal:

**Equation 7:**

$$T_{anterior} = T_{posterior}$$

Combining (6) and (7) we obtain the following equation:

**Equation 8:**

$$v_{zip} + v_{unzip} = \frac{(S_{anterior} - S_{posterior})}{\eta}$$

Since $v_{zip} > 0$ and $v_{unzip} > 0$, therefore,

$$S_{anterior} - S_{posterior} > 0, or\ S_{anterior} > S_{posterior}$$

The model, therefore, predicts the existence of differential adhesion which would result in forces driving adhesion to the anterior neighborhood.
